# Two-colour Imaging Fluorescence Correlation Spectroscopy (ImFCS) probes the influence of the juxta-membrane actin-cortex

**DOI:** 10.64898/2025.12.24.696335

**Authors:** Sankarshan Talluri, Pooja S Krishna, Thorsten Wohland, Satyajit Mayor

## Abstract

Here we extend Imaging Fluorescence Correlation Spectroscopy (ImFCS), together with bright, photostable fluorophores, to quantify how the juxta-membranous actin cortex influences plasma membrane dynamics. This problem is technically challenging because the membrane is only a few nanometres thick, whereas the axial resolution of live-cell fluorescence imaging is on the order of 100 nm. We address this by engineering a membrane-anchored actin reporter that combines a weakly interacting actin-binding domain with bright, photostable fluorophores optimized for two-colour ImFCS. This design enables the generation of simultaneous, co-registered diffusion maps where the mobility of the reporter serves as a dynamic sensor for the local density of the cortical actin network. The actin-binding reporter displays diffusion coefficients spanning more than two orders of magnitude, in stark contrast to an otherwise similar reporter that cannot bind actin, which diffuses much faster and within a much narrower range. Thus, the mobility of the reporter becomes a sensitive spatial readout of actin-mediated constraints. By comparing, on a pixel-by-pixel basis, the diffusion of the actin-binding reporter with that of the non-binding mutant, we construct an actin-occupancy map that quantifies the fraction of the membrane functionally influenced by cortical actin. Using this approach, we find that the juxta-membrane actin cortex modulates transport over more than 80% of the cell surface. Our two-channel ImFCS strategy establishes diffusion-based comparison as a general methodology to map the functional footprint of the cortical cytoskeleton on live cell membranes.

## INTRODUCTION

Imaging Fluorescence Correlation Spectroscopy (ImFCS) is a variation of conventional point-based FCS. Instead of computing the auto-correlation function of fluorescence fluctuation in a confined volume (e.g. diffraction limited volume of focused laser illumination), ImFCS computes correlation functions from pixels imaged over a large area, thereby generating spatial maps of the autocorrelation of fluorescence fluctuations (1, 2). By fitting each autocorrelation function to a model that adequately accounts for the fluctuations (e.g. 2D diffusion of a molecule in a membrane), a spatially resolved interrogation of the heterogeneity of the landscape that influences these fluctuations may be described.

The simultaneous measurement of fluctuations from two different molecules in an FCS measurement provides information about local properties of the underlying medium such as the relative mobilities of molecules or the binding interactions of the two species in the same location in the plasma membrane of a cell. Fluorescence Cross Correlation Spectroscopy (FCCS) is one such two channel measurement which has been successfully implemented in live cells. Here most applications have focussed on the cross-correlation amplitude to measure binding rather than generating independent diffusion coefficient maps (1, 3). This is because spectral bleed through between channels can complicate quantitative interpretation of the diffusion coefficient. This can only be resolved by technically involved strategies such using pulsed interleaved excitation strategies or fluorescence lifetime gated correlation spectroscopy to mitigate cross emission talk (4, 5). In the context of the plasma membrane, these limitations have meant that dual channel fluctuation measurements are relatively rare, and have not been used to construct two independent, co-registered diffusion maps that can be compared at the pixel level across an entire cell surface.

As described in this study, diffusion maps of the cell surface provide information about the underlying cortical actin cytoskeleton that supports the plasma membrane in animal cells. A dense, juxta-membrane actin network shapes cell morphology, supports membrane mechanics and constrains the lateral mobility of lipids and proteins, thereby influencing receptor clustering, signalling reactions and endocytic traffic, and is a central regulator of plasma membrane organisation and function (6–10). Models such as the anchored picket fence propose that actin-anchored transmembrane proteins form semi-permeable boundaries that compartmentalise diffusion into corrals and promote hop diffusion of membrane components (11–13). More recent work has shown that membrane proximal F-actin (MPA) can integrate upstream signalling inputs to locally restrict protrusions and direct cell migration, highlighting MPA density as a key structural parameter in cell polarity (14–17).

Visualizing the actin-cortex by electron and fluorescence microscopy reveals that the membrane is draped across this sub-structure (12, 18, 19). Since the plasma membrane is about 5 nm thick and the best axial resolution for live cell imaging is total internal reflection fluorescence (TIRF) microscopy where the evanescent field integrates signal from a depth of ∼100 nm, the fluorescence intensity of any actin probe is an axial average that conflates actin filaments that are close enough to modulate molecules at the membrane with those that are probably further away and may not have a direct effect. Since an actin binder would need to physically interact with the actin to be slowed down by it (20), diffusion measurements of membrane anchored actin binders using ImFCS presents a unique solution to the limited resolution of intensity-based measurement. By being able to image the cortical-actin cytoskeleton, and a membrane associated molecule at the same time (20–22), in distinct, spectrally resolved channels, we aimed to use the property of ImFCS to convert the spatial distribution of the mobility of a membrane anchored actin binder into a spatially resolved map of actin influence.

In this work, motivated by the need for a readout of the juxta-membrane actin cortex, we developed a minimally perturbing, high signal to noise actin reporter that is explicitly optimised for two channel ImFCS. The probe is based on coupling a highly bright and photo-stable fluorophore (mStayGold or Halo-tagged with JF657) to an actin-binding domain derived from F-Tractin (23). The low affinity of the reporter allows it to dynamically bind and unbind to an actin network that is also remodelling at timescales slower than the diffusion of the binder on the membrane, while high brightness of the actin binder permits reliable correlation analysis over a wide range of diffusion regimes. As a non-interacting membrane tracer, we used a mutation of the actin-binding domain that abrogates association with F-actin.

We measured the diffusion of the actin binder along with the mutated non-binder that serves as a passive membrane marker in the second channel, enabling us to obtain simultaneous, co-registered diffusion maps for the binder and the tracer in every pixel of the field of view. By comparing the local mobility of the actin reporter to that of the non-binding tracer in the same region, we compute a differential diffusion map that reflects the effective occupancy of membrane proximal actin. This enables a new way to quantify the area fraction of membrane that is functionally influenced by actin. This analysis suggests that actin retards diffusion over a large majority of the cell surface. In the present paper, we focus on the development, characterisation and implementation of this weak binder based, two channel ImFCS approach. We describe the design of the actin reporter, the optical and analytical pipeline required to extract robust two channel diffusion maps, and the basic features of the resulting actin and tracer diffusion landscapes, with an emphasis on the methodological advances that make this form of juxta-membranous mapping possible.

## MATERIALS AND METHODS

### Plasmids and Constructs

Four custom constructs were generated for this study by Twist Biosciences using the pTwist CMV Puro vector. All constructs were modified from the MPAct probe (14) retaining the CaaX membrane-targeting motif. The fluorophore was replaced with either mStayGold (24) or a HaloTag. The actin-binding domain was replaced with either F-Tractin-Opt (23), referred to as TractO, or its filament-binding mutant, F-Tractin-Opt-FA (23), referred to as TractO*. This resulted in four variations of the membrane binding probe: CaaX-StayGold-TractO, CaaX-StayGold-TractO* (containing the FA mutant), CaaX-HALO-TractO, and CaaX-HALO-TractO*.

### Cell Culture and Labelling Protocols

CHO-K1 cells were seeded onto Helmanex-cleaned glass-bottom dishes. At 48 h post-seeding, cells were co-transfected with plasmid constructs using JetPrime transfection reagent (Polyplus) according to the manufacturer’s protocol. Each transfection was performed with 1 µg of each construct. The following plasmid combinations were tested in separate experiments:

1. CaaX-HALO-TractO + CaaX-mStayGold-TractO
2. CaaX-HALO-TractO*+ CaaX-mStayGold-TractO
3. CaaX-HALO-TractO*+ CaaX-StayGold-TractO*

After four hours of transfection, cells were washed with complete medium and incubated with 40 nM JF657-Halo dye (From Janelia Materials) for 30 min at 37 °C. Excess dye was removed by washing twice with complete medium, with each wash followed by incubation on a rocker for 15 min at room temperature. For all plasmid combinations, imaging was performed in a 1× M1G buffer supplemented with 5 mg/mL BSA.

### Implementation of ImFCS on a TIRF microscope and its analysis

Cells were imaged in TIRF mode maintained at room temperature of 20 degrees on an inverted epifluorescence microscope (Ti-E, Nikon) equipped with a objective (100×, NA 1.49, Nikon). Excitation used 488 nm and 640 nm laser lines (Agilent, MLC300) combined through the Nikon semi-automatic TIRF illuminator. The back focal plane was illuminated using a 405/488/561/647 dichroic (Chroma). Emission passed the same objective and dichroic and was split with a 600dcxp long pass filter on the Cairn MultiCam, and cleaned up with a 525/50 filter (488 channel) and a 685/50 filter (640 channel), before detection on 2 orthogonally placed, sCMOS (Prime95B) cameras. For ImFCS, the camera ROI was operated in Sensitivity mode with an ROI of 300×95 pixels, with the clear mode set to Pre-sequence. An external trigger from a pulse generator was used to fire both the cameras simultaneously and the cameras were operated in Trigger First mode that runs the acquisition on the cameras’ internal timer after a synchronized starting pulse. Alignment between two cameras was made manually using a 100nm tetraspeck bead slide, before each experiment. Data were acquired with Micromanager 2.0 subsequent analysis used Fiji and the ImFCS plugin (v1.613) (25). Laser power at the back focal plane was set at 3 mW in the 488 channel and 2.5mW is the 640 channel. Unless specified, the physical pixel size on the sCMOS was 11µm, was magnified a 100 times to get a 110nm pixel size in the image. Typically 50,000-70,000 frames were acquired at ∼ 2 ms per frame.

Image stacks were imported into Fiji and processed with ImFCS 1.613. Unless noted, analysis used 3×3 binning, objective NA 1.49, refractive index matched to the medium, and emission wavelengths λ = 525 nm (488 channel) and λ = 685 nm (640 channel). Photobleaching correction was applied by fitting and removing a fourth-order polynomial with a sliding window of 1,000 frames in the plugin’s polynomial bleach corrector.

Autocorrelation curves were computed per pixel using ImFCS with correlator parameters suited to the frame rate and field of view. Typical settings were p-q = 16-10 for cell membranes. For each pixel the plugin produced a 2D autocorrelation function for fitting.

Autocorrelation functions were fitted with a two-dimensional one-component diffusion model appropriate for TIRF sampling (26).

## RESULTS AND DISCUSSION

### Acquisition of simultaneous, two channel, spatial diffusion maps

To enable a local, mobility-based readout of the juxta-membrane actin influence, we implemented a two-channel Imaging Fluorescence Correlation Spectroscopy (ImFCS) workflow in total internal reflection fluorescence (TIRF) microscopy mode (schematic in **Figure 1a**). Our design builds on previous work (2, 20), with two primary modifications. Firstly, we generate diffusion maps in two channels at the same time by combining the use of fluorescent proteins and organic dyes with well separated emission spectra (**Supplementary Figure S1a**). Secondly, to generate a useful functional map of diffusion of the cell membrane, we acquire a high signal to noise signal for each correlation curve at every pixel on the membrane. The Signal to Noise Ratio in FCS experiments is given by the following expression (27), where

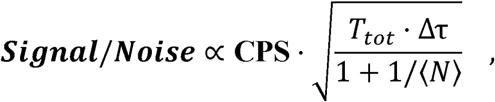

**Figure 1.**
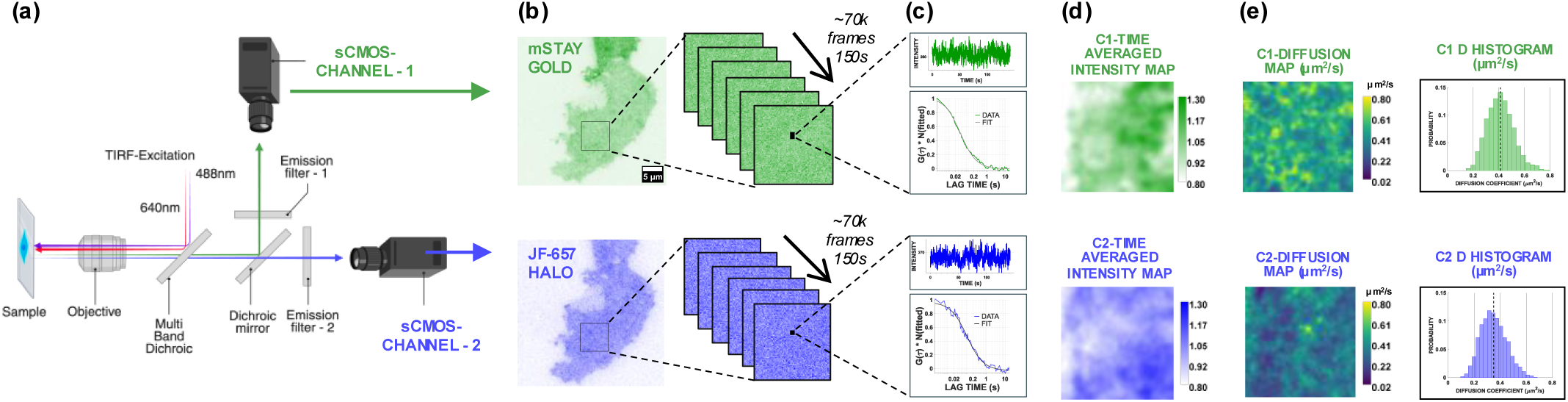
Two channel imaging FCS. **a)** Schematic of the imaging configuration used for the experiments. **b)** Image of the same cell in channel 1 (488nm excitation, labelled with mStayGold) and channel 2 (640nm excitation, labelled with JF-657 HALO tag). The highlighted region is selected for the ImagingFCS measurement, from which ∼70k frames are acquired. **c)** The intensity trace from each pixel in the ROI is autocorrelated to generate an autocorrelation curve that is fitted to extract the diffusion coefficient in that pixel. An independent diffusion coefficient is extracted for each channel. A representative intensity trace from one pixel is shown (channel 1 – green, channel 2 – blue), followed by the calculated autocorrelation function (channel 1 – green, channel 2 - blue) and the fit (black). **d)** The time averaged intensity map for each ROI is generated by summing all the intensities at all time points and then is normalized to the mean intensity of each channel. Values on this map correspond to the average density of labelled molecules in the duration of the acquisition and can be thought of as a space-time density map. **e)** Diffusion coefficient map for each channel, obtained by fitting autocorrelation curves for each pixel. Histogram shows the probability distribution of the fitted diffusion coefficient values shown in the map.

**CPS** is the Counts per Particle per Second which is essentially the brightness of the particle. **T_tot_** is the total duration of the acquisition, Δτ is the time interval of one frame and **N** is the number of particles in the observation volume. CPS is the only parameter among these that influence the S/N ratio linearly making it the most important parameter influencing the precision of FCS measurements (28). CPS is influenced by intrinsic fluorophore brightness (quantum yield times extinction coefficient) and the laser power used for fluorescence activation. Fluorophore quantum yields are limited by the properties of the fluorophore and cannot be changed, whereas the laser power can be increased to increase photon yield. However, conventional fluorescent proteins are modest in brightness which reduces the signal to noise of correlation functions. A further concern in FCS measurements where binding and diffusion occur within the observation volume is cryptic photobleaching. In this context, fluorophores can bleach while a molecule remains inside the illuminated volume (27). Because bound or slowly moving molecules have longer residence times, they are disproportionately likely to bleach, biasing the analysis against long-lived states. The practical consequence is that slow or bound populations appear artificially short-lived, which compresses the apparent separation between “bound” and “free” mobilities and can mask long-lived interactions. This effect has also been experimentally seen for membrane associated actin binders (20) and transcription factor binding in the nucleus (29). Therefore, an ideal fluorophore for FCS (and most other single molecule measurements) is one that is bright (high quantum yield and extinction coefficient) and photostable (resistant to high laser powers). To address both the considerations described above, we designed an experimental system with a combination of a fluorescent protein – mStayGold (24), and an organic dye – JF-657 (30). mStayGold and JF-657 are two fluorophores with best-in-class photostability and brightness. Point-FCS with mStayGold has been very promising and has led to high signal to noise, photobleaching free, and massively parallel FCS measurements of diffusion and binding constant measurements (31). JF-657 is a cell permeable analogue of AttoFluor 647N developed by the Lavis laboratory with the highest photostability among all the far-red excitation dyes that they tested (30).

Labelled and transfected cells are excited simultaneously with 488nm and 640nm laser line, and emission from both the fluorophores is split by using a dichroic mirror, and each channel is collected on orthogonally placed sCMOS detectors that were co-registered using fiducial bead images acquired in the same optical configuration. The details of the filters and the emission spectra of the fluorophores are described in **Supplementary Figure S1a** and in the **Methods Section**.

From each acquisition, we generated both intensity (**Figure 1d**) and diffusion (**Figure 1e**) maps. The acquisition consists of a time series of TIRF images, from which the temporal intensity trace at each pixel is used to compute a pixel-wise autocorrelation function (ACF; **Figure 1c**). The intensity maps are generated by taking a sum projection along the acquisition time and are normalized to the mean intensity of the sum projection (**Figure 1d**). After photobleaching correction, each ACF is fit to a two-dimensional diffusion model to extract a diffusion coefficient D per pixel (**Figure 1c**). The resulting diffusion maps (**Figure 1e**) are therefore derived from the same raw frames as the intensity maps but encode dynamic information that is not available from diffraction-limited intensity maps. This paired representation provides a direct way to compare the actin-binder mobility with the density of the membrane molecule of interest in the same optical pixel.

### Weak actin-binder provides a mobility-based reporter of the juxta-membrane actin cortex

It has been previously shown that the dynamics of the membrane proximal actin are different from the dynamics of the actin in the entire cellular pool (14). To map the membrane proximal actin distribution and dynamics we modified an existing membrane associated actin-binder (14), replacing mEGFP/RFP with mStayGold/Halo-tag for spectrally separated visualization. The construct (**Figure 2a**) binds to the membrane with a CaaX domain taken from K-Ras which consists of a site for the addition of a single farnesyl moiety and has a polybasic region that interacts with anionic lipids in the membrane. This is sufficient to localize the associated protein to the plasma membrane. As a reporter of local actin concentration we used an optimized F-Tractin_opt_ construct (referred to TractO) and a point mutant F29A F-Tractin_opt_ (TractO*) that results in loss of actin binding as a non actin-binding control protein (23). TractO is a 21 amino acid truncated F-Tractin peptide that binds to actin with less affinity than F-tractin, and has reduced actin bundling capacity compared to F-Tractin (23). These are important characteristics in any actin-reporter. Both these constructs were expressed in CHO-K1 cells transiently and visualized by TIRF microscopy.

**Figure 2.**
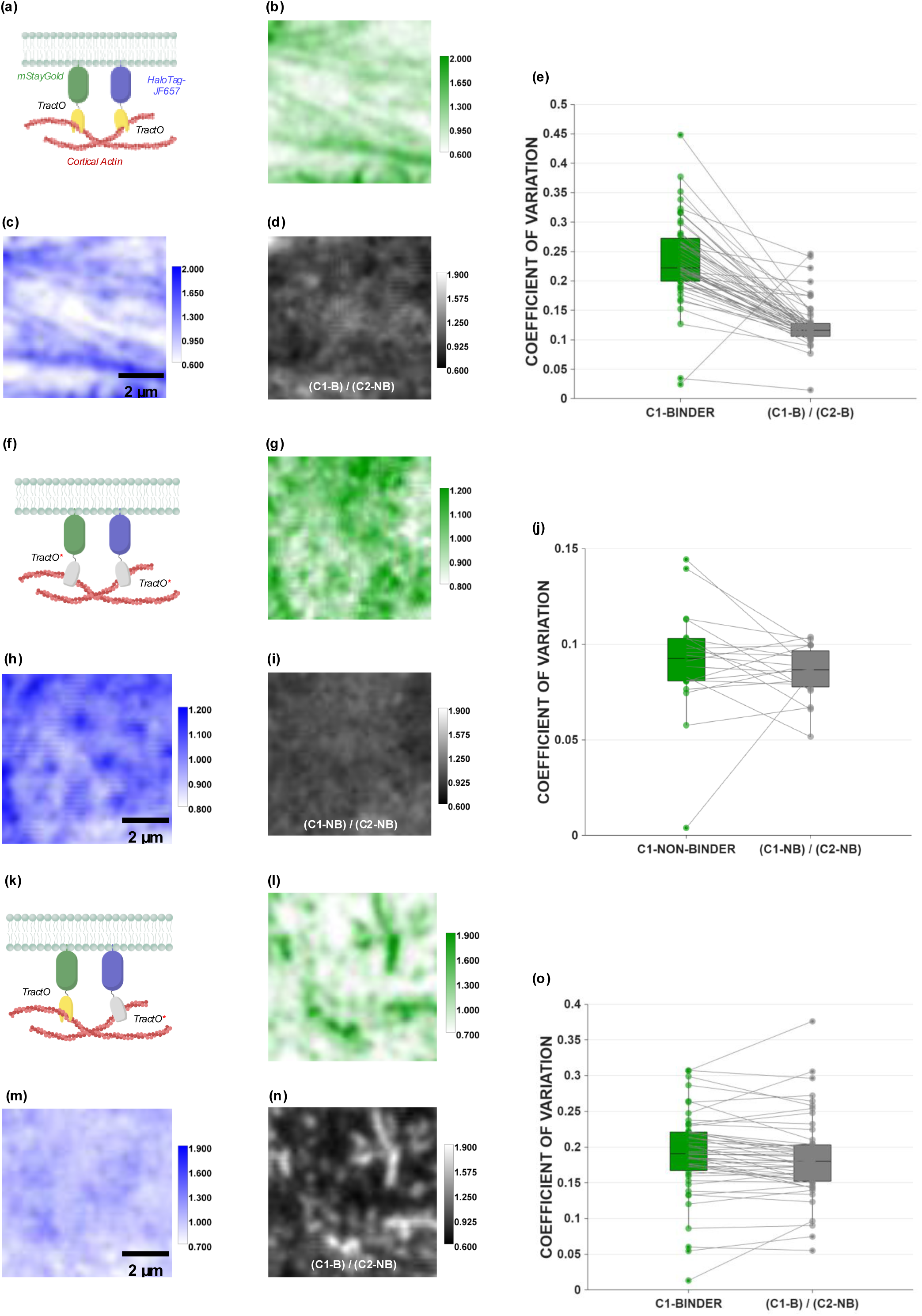
Time averaged actin binder intensity maps. **a)** Schematic of the membrane anchored actin binders depicts the two proteins inserted into the inner leaflet of the membrane bilayer with a CaaX domain, linked to with mStayGold/Halo-Tag and an actin-binding domain, TractO. **b,c)** The time averaged intensity maps for channel 1 mStayGold-TractO (b) and channel 2 Halo-Tag-JF657-TractO (c) are each normalized to their respective means. LUT shows the calibration bars for each channel. **d)** Intensity ratio map of the Channel 1 divided by the Channel 2 intensity maps. **e)** Pairwise comparison of the coefficient of variation calculated for all the ROIs imaged. The CV of the C1 channel (in b) is compared with the CV of the corresponding ratio image (in d) (pooled data from 30 ROIs of ∼50µm2, taken from 18 cells). **f)** Schematic of the membrane anchored actin non-binders depicts membrane associated CaaX domain, labelled with mStayGold/Halo-Tag and the mutated actin binding domain, TractO*. **g, h)** The time averaged intensity maps for channel 1 (g) and channel 2 (h) are each normalized to their respective means and shown along with their respective LUTs. **i)** Intensity ratio map of channel 1 divided by channel 2. **h)** Pairwise comparison of the CV of the C1 channel (g) compared to the CV of the corresponding ratio image (i) from data derived from 20 ROIs of ∼50µm2, taken from 13 cells. **k)** Schematic of the membrane anchored actin binder, mStayGold-TractO and non-binder, HaloTag-JF657-TractO* expressed in the same cells. **l,m)** The time averaged intensity maps for channel 1 mStayGold-TractO (l) and channel 2 Halo-Tag-JF657-TractO* (m) normalized to their respective means are shown along with along with LUTs. **n)** Intensity ratio map, of the channel 1 (l) divided by channel 2 (m) intensity maps. **o)** Pairwise comparison of the CV of the C1 channel (l) is compared with the CV of the corresponding ratio image (n), from data derived from 51 ROIs of ∼50µm2, taken from 30 cells.

The membrane linked actin-binder provides a way to visualize juxta-membrane actin structures in terms of higher intensity features of the time-averaged intensity map. These high intensity features in the map have at least two contributions, one from the intensity that may be attributed to actin-architecture proximal to the membrane impeding diffusion of the binder, and another from other membrane features (membrane protrusions, curvature, and others) that would also be reflected in the time-averaged intensity maps. To resolve this further, we examined the time-averaged intensity distribution of the actin-binder, TractO, and that of the actin-non binder, TractO*, in different combinations. When expressed in the same cell, while mStayGold-TractO and Halo-JF647-TractO exhibit very similar patterns (**Figure 2b,c**), mStayGold-TractO* and Halo-JF647-TractO* exhibit almost uniform distributions (**Figure 2g,h**). To normalize the features exhibited by a probe in one channel we built normalized ratio intensity maps by dividing the normalized intensity in one channel by the other in the corresponding channel. As noted by the graphs of the Coefficient of Variation (CV) of the intensity distribution in the TractO channel, the CV only reduces when the intensity maps of the TractO is normalized to intensity in the corresponding TractO channel (**Figure 2e**), and not when it is normalized by the TractO* channel (**Figure 2o**). These observations suggest that a significant set of features as outlined by the intensity maps of actin-binder probe, TractO, are independent of other features from the membrane as described by the TractO* map. Therefore, we maintain that the TractO intensity distribution provides an approximation of the juxta-membrane actin distribution. By constructing movies of the distribution of the actin-intensity images (**Supplementary Movie 1; and montage in Supplementary Figure S2**), we note that this actin distribution consists of very slowly varying component (>100sec) as well as a dynamic component that appears to remodel on the 10’s of seconds time scale. The slowly varying component could represent stress fibers that are membrane proximal, and the very dynamic actin-structures are likely the contractile cortical actin meshwork. However while the high intensity features visualized by the TractO probes outline an imprint of the MPF, it is unclear if this is represents the full spatial extent of MPF. To address this question we have turned to ImFCS using these fluorophores.

One reason we are able to visualize these features at this temporal resolution, is because these fluorophores are remarkably photostable; they bleach to only 20-30% of their starting value after ∼2.5 minutes of illumination at lasers powers of ∼2mW at the back focal plane (**Supplementary Fig. S1b; Methods**). For comparison the normalized intensity trace of an mEGFP labelled membrane protein is almost completely bleached at half the laser power used for imaging mStayGold. These features will also allow long term ImFCS measurements to capture full time scale of the dynamics of the membrane proteins as well as the underlying changes in actin architecture. Average diffusion coefficients for a given ROI (yellow box, **Figure 3a,b**) are shown for each channel in **Figure 3c,e**. In both channels TractO diffusion coefficient is slowed down about 5-6 fold compared to average diffusion coefficient TractO* in the same pixels. This is a large shift in mobilities compared to other studies (20), where a difference of less than two fold is observed between actin-binders and non-binders (20, 32, 33). This is likely because of the high photostability of the fluorophores used in our studies, minimizing cryptic photobleaching in the measurement that may lead to artificially inflated mobility estimates. Diffusion coefficient distributions for the binder and non-binder, measured independently in both spectral channels exhibit a range of values (**Figure 3d,f**). TractO exhibits a range spanning more than two orders of magnitude (filled bars in **Figure 3d,f**, Coefficient of Variation (CV) C1-binder = 0.593, C2-binder = 0.5372). In contrast, the non-binder, TractO* exhibits a smaller range within an order of magnitude (CV of C1-non-binder = 0.323, CV of C2-non-binder = 0.3525) and has a more right shifted diffusion coefficient distribution (hollow bars, **Figure 3d,f**). Thus, the actin-binding probe distinguishes actin influenced membrane environments not only by a change in average mobility, but also by the increased spread of the diffusion coefficient distribution. This widening is significant because it implies that the actin influence experienced by the probe is not spatially uniform. Instead, the binder (TractO) samples a heterogeneous mobility landscape, with pixels occupying distinct transport characteristics. This raises an important question. Does the increased CV reflect heterogeneity in the juxta-membrane actin influence, or does it mostly reflect technical variability? To answer this, we tested whether the spatial heterogeneity of diffusion covaries with actin architecture and whether it is reproduced across channels.

**Figure 3.**
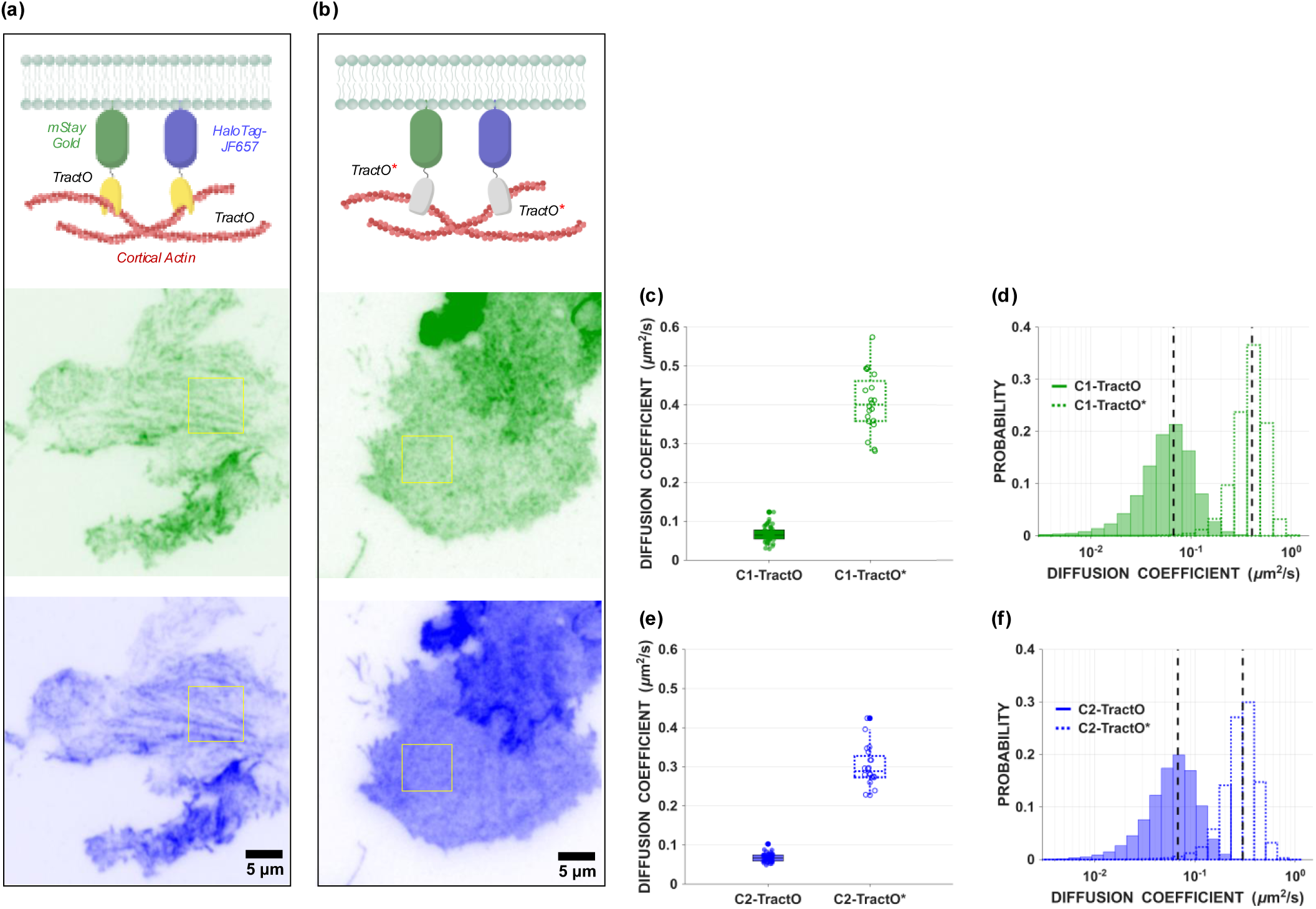
Diffusion measurements of TractO and TractO*. **a,b)** Schematic of mStayGold and HaloTag-JF657 labelled TractO (a) and TractO* (b) expressed in cells and representative TIRF images of the same cell expressing mStayGold (green) and HaloTag-JF657 (blue) constructs of the same protein. The boxed region (yellow) is selected for ImFCS. **c,e)** Average diffusion coefficients across all the pixels of the TractO in channel 1 (green) and TractO* in channel 2 (blue) in the ∼50µm^2^ region selected for ImFCS. Data points represent the diffusion coefficient of the binder - TractO (filled circles) taken from 30 ROIs of ∼50µm^2^ from 18 cells and the non-binder - TractO* (open circles) from 20 ROIs of ∼50µm^2^ from 13 cells. **d,f)** Histogram depicts the probability distribution of diffusion coefficients determined from all the ROIs for the channel 1 (green) and channel 2 (blue), TractO (filled bars) and TractO* constructs.

### Two-channel diffusion maps reveal actin architecture

If the increased intensity of the TractO in the image is because of a higher actin concentration in that pixel, the expectation is that the diffusion in each pixel will be negatively correlated to the time averaged TractO intensity in that pixel. Visually this negative correlation between these pixels can be observed in the intensity (**Figure 4b,f**) and diffusion maps (**Figure 4c,g**) and is shown graphically in **Figure 4i**. To understand how the spatial maps are correlated across channels, we examined the correlation between the TractO intensity in channel 1 (mStayGold) and TractO diffusion in channel 2 (HaloTagJF657). Here as well, a similar trend between the mStayGoldTractO intensity (C1-Intensity) and HaloTagJF657TractO diffusion (C2-Diffusion) in the other channel is observed (**Figure 4j**).

**Figure 4.**
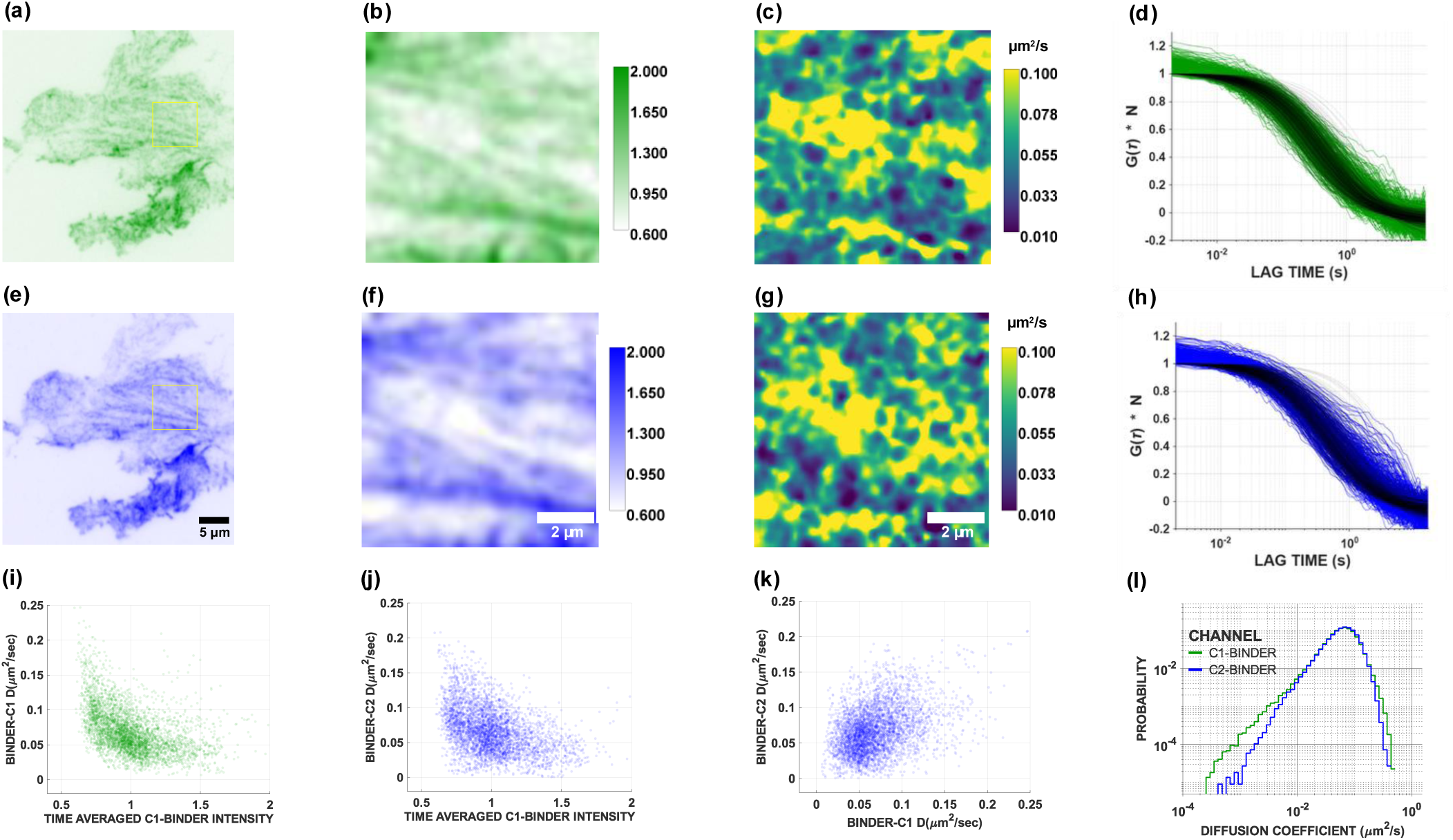
Spatial map showing the effect of TractO density on its diffusion coefficient. **a-h)** Representative images of mStayGold-TractO in channel 1 (a,b) and HaloTag-JF657-TractO in channel 2 (e,f). The square overlay represents the region used for the ImFCS measurements (b,f). Time averaged intensity maps for channel 1 (b) and channel 2 (f) are each normalized to their respective mean intensity values and shown along with respective LUTs. Diffusion maps (c,g; and their corresponding LUT) are derived from fitting the autocorrelation curves for each pixel (d, h) shown with their corresponding fits. The computed correlation curves are in green (channel 1) and blue (channel 2) and the fits are shown in black. The fits and the raw data are normalized to the inverse of the fitted y-intercept. **i, j)** Scatter plots depict time-averaged intensity of each pixel in channel 1 derived from intensity images in panel b, plotted against the corresponding pixel value of the diffusion of the binder in channel 1 (i) and channel 2 (j). **k)** Diffusion coefficient of each pixel in the channel 1 D map (c) plotted against the corresponding pixel in channel 2 (g). **l)** Probability distributions of the fitted diffusion coefficient values in panels c and g.

We next examined if the actin binder diffusion maps in both channels (**Figure 4e,h**) are covarying. As observed in the scatter plot (**Figure 4k**) comparing each pixel in the diffusion map the positive correlation is unmistakable. The correlations shown in **Figure 4i-l** serve as an important positive control for the measurement and analysis pipeline.

Although the correlation is evident, it is not advisable to parametrize such a non-linear dependence with traditional measures of linear correlation like PCC. Additionally, the diffusion coefficient distributions measured here are average values and have a contribution from stochastic noise in the estimation of the diffusion coefficient and another contribution from actual biological heterogeneity. It is not trivial to separate the two contributions from each other and the presence of the noise will have an impact on the value of the correlation measured as one would only expect to detect correlations in the signal and the noise will remain uncorrelated. For these reasons we have used another correlation index called the Threshold Overlap Score that is more suited to such datasets.

The Threshold Overlap Score (TOS) (34) is a metric designed to quantify the localization of two signals in microscopy images by measuring their overlap relative to chance. The calculation first measures the observed fractional area of overlap, which is the fraction of pixels that are present in a defined threshold for both signal 1 and signal 2. This observed overlap is then compared to the overlap that would be expected purely by chance, assuming a null distribution where the signals are uniformly and independently distributed. Finally, this comparison is linearly rescaled to a standardized score ranging from **-1** (indicating maximum anti-colocalization, or less overlap than chance) to **+1** (indicating maximum colocalization, or more overlap than chance), with a score of **0** representing non-colocalization, where the observed overlap is the same as expected by chance.

We have used this metric to generate a TOS matrix, which systematically calculates the TOS value at all combinations of the thresholds specified. Modifying it from the way it was used in the paper that first described the metric (34), we divided both the signals that were being compared into deciles (from data such as depicted in **Figure 5a,c**) and compared the Threshold Overlap Score for each combination, which then populates the TOS matrix (**Figure 5b,d**). The utility of this matrix-based approach is its ability to characterize localization patterns across all signal ranges – regions with high diffusion vs regions with low diffusion. This allows it to identify and distinguish mixed localization patterns within a single sample—for instance, revealing strong colocalization only at certain thresholds (on-target signals) while simultaneously showing non-colocalization at the rest of the thresholds (background signals). Additionally, the key feature of the TOS metric is that it does not assume a linear correlation in signal intensities, making it more broadly applicable. This provides a more detailed and robust characterization of localization than is possible with traditional, single-value metrics. This analysis provides flexibility to design the thresholds according to specific structure of each dataset. In this case, the regions of high diffusion/intensity or low diffusion/intensity represented regions where the expected effect of the noise in the estimation would be the weakest and the contribution of the signal to be dominant. **Figure 5b** shows the TOS matrix calculated for the scatter plot in **Figure 5a**, representing the relationship between the binder diffusion in channel 2 with the binder intensity in channel 1. Each square grid in the heat map is the TOS value according to the definition in the source paper (34) at that combination of threshold values. From the scatter we can see that points with a high binder diffusion coefficient seem to be more preferentially present at lower binder intensities, this implies that there is a lower likelihood of finding points with high diffusion coefficients at high binder intensities. This lower likelihood is captured by a negative TOS score in the top left corner of the TOS matrix (**Figure 5b**). Similarly, the negative TOS score can be observed in the bottom right corner of the matrix as well showing that points with low diffusion coefficients and low binder intensities co-occur less frequently than expected by chance. A singular TOS score was defined as the average of the points contained in the dotted squares in the top left and bottom right of the matrix.

**Figure 5.**
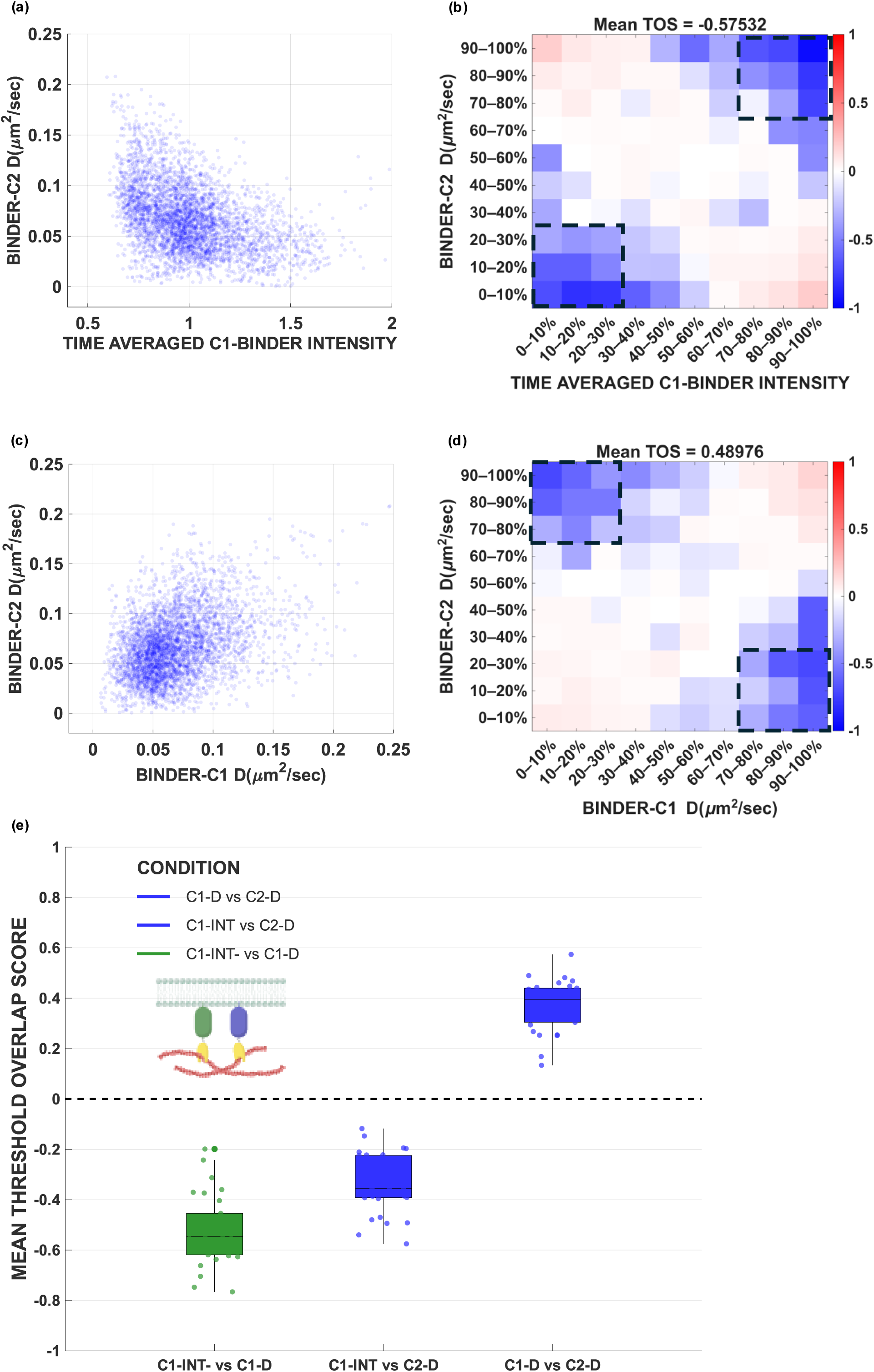
Threshold overlap score matrix as a tool to parametrize colocalization. **a)** Intensity of each pixel in the channel 1 (mStayGold-TractO) plotted against the corresponding pixel value of the diffusion of in channel 2 (HaloTag-JF657-TractO). **b)** TOS matrix for the scatter plot in panel a where the Mean TOS value (average of the highlighted 18 squares), is displayed on the top of the Matrix. **c)** Diffusion coefficient of each pixel in channel 1 (mStayGold-TractO) plotted against the corresponding pixel value of the diffusion in channel 2 (HaloTag-JF657-TractO). **d)** TOS matrix for the scatter plot in panel c. **e)** Box plot highlighting mean TOS matrix scores as indicated. Data pooled from 30 ROIs of ∼50µm2, taken from 18 cells.

It is to be noted that the particular fashion in which the matrix has been defined, a negative TOS score does not necessarily imply a negative correlation in the distributions being compared. A negative TOS score simply indicates that the overlap is less than what is expected by chance. The values of the deciles being compared also need to be considered to assign a sign to the final correlation. A negative TOS between the high-high and low-low deciles of the distributions is explained by an overall negative correlation between the variables being compared. However, in the case of **Figure 5c**, where the data is positively correlated (channel 1 binder diffusion vs channel 2 binder diffusion) we expect to see a negative overlap between the high-low and low-high deciles. This flip in the location of the negative TOS values can be seen in TOS matrix in **Figure 5d**. This implies that the lower deciles of one distribution do not overlap with the higher deciles of the other distribution, a scenario only possible in a positive correlation. In the analysis, the sign of the negative TOS values taken from the top left and bottom right of the matrix is changed as they indicate a positive correlation (dotted lines in **Figure 5d**) and the negative TOS values from the top right and bottom left of the matrix are left as they are because this combination measures the overall negative correlation in the variables.

Pooling data from multiple cells together per condition in **Figure 5e**, with binders expressed in both channels, there is a strong negative correlation between the channel 1 binder intensity and channel 1 binder diffusion (green box plot, **Figure 5e**), and a slightly lower but strong negative correlation between the channel 1 binder intensity and channel 2 binder diffusion. There is also a strong positive correlation between the diffusion of both binders. This is consistent with the trends in the images that were visible qualitatively in **Figure 4i-k**.

### Diffusion-difference mapping yields a functional actin occupancy metric

We asked what fraction of the pixels experience the influence of the underlying actin by examining the diffusion of the membrane anchored TractO on the membrane and the non-binder, TractO* in the same pixel. When expressed together in the same cell (**Figure 6a**), the average diffusion of the TractO* in each ROI was higher than that of the TractO (**Figure 6b**), as expected. This implies that the actin-binding is responsible for slowing down the average diffusion in each ROI. Moving beyond average values, we used the availability of the diffusion coefficient information of the non-binder and the binder in the same pixel to examine the spatial influence of actin on the diffusion of membrane components.

**Figure 6.**
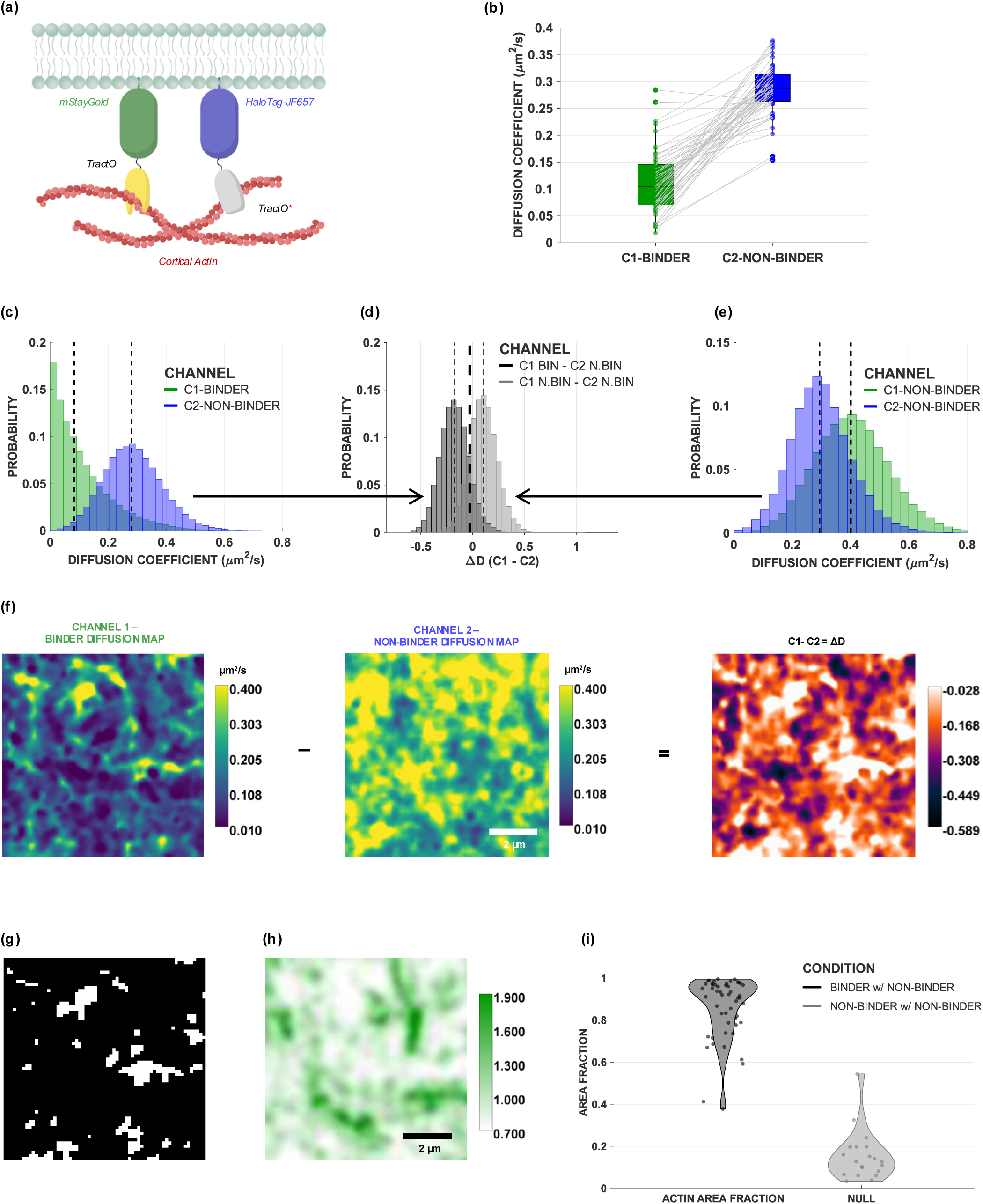
Measuring area fraction of actin coverage at the cell membrane. **a)** Schematic of the constructs used to determine area coverage of actin at the membrane. **b)** Box plot showing the differences in average diffusion coefficient of the mStayGold-TractO (green) and HaloTag-JF657-TractO* (blue) in measurements of each individual region. Paired regions are connected by a gray line in data taken from 51 ROIs of ∼50µm2, taken from 30 cells. **c-e)** Probability distribution of diffusion coefficients determined from corresponding pixels in cells simultaneously expressing mStayGold-TractO and HaloTag-JF657-TractO* (c; data from 51 ROIs taken from 30 cells) or mStayGold-TractO* and HaloTag-JF657-TractO* (e; data from 20 ROIs of ∼50 µm2, taken from 13 cells). Histograms (d) show the difference of two diffusion coefficient distributions; C1-Binder (mStayGold-TractO) — C2-Non-Binder (HaloTag-JF657-TractO*) dark grey, C1-Non-Binder (mStayGold-TractO*) and — C2-Non-Binder (HaloTag-JF657-TractO*), light grey. Thick dashed line represents threshold chosen to separate both distributions (mean minus one S.D. of null (light grey) distribution. **f)** Representative diffusion maps of the mStayGold-TractO in channel 1 and the HaloTag-JF657-TractO* in channel 2 and their difference map (in the blue orange LUT). **g, h)** Thresholded difference map (g) where all the pixels lesser than the threshold of −0.028 (in panel f) have been set as black, representing regions where the effect of the actin on the binder is distinctly detectable along with corresponding time-averaged intensity map of mStayGold-TractO (h). **i)** Violin Plot depicts the area fraction of the region of the membrane that exhibits a slowed diffusion of the indicated membrane protein from the estimation made with the same threshold (−0.028) for the null distribution (light grey). Data derived from panels (c) and (e).

To define a threshold to classify pixels exhibiting a slower diffusion of TractO versus TractO* we used the difference of TractO versus TractO* diffusion coefficient distributions in each pixel (**Figure 6c**). The difference in diffusion coefficient of TractO* in the two channels was used to generate a null distribution representing the differences in diffusion coefficients of the non-binders expected without any actin involvement (light grey, **Figure 6d**). TractO* in channel 2 (HaloTag-JF657-TractO*) has a diffusion coefficient similar to the TractO* in channel 1 (mStayGold-TractO*); they differ only by 0.1µm^2^/s (**Figure 6e**). Similarly, the difference of the diffusion of TractO (mStayGold-TractO) versus TractO* (HaloTag-JF657-TractO*) were utilized to generate the histogram of the pixelwise slowdown of the TractO as compared to TractO* (dark grey, **Figure 6d**). This operation is visualized in **Figure 6f**, where the TractO diffusion coefficient map is subtracted from the TractO* diffusion coefficient at each pixel to generate the difference map (rightmost panel). The final difference map is shows the relative reduction in the diffusion coefficient of the binder (TractO) in each region as compared to that of the non-binder (TractO*). By selecting a threshold at the value of one standard deviation less than the mean of the null distribution (thick, dashed line in **Figure 6d**, Δ D = −0.028), and applying this threshold on the difference image, we find that most of the pixels (∼80%) are under this threshold (black regions in **Figure 6g**). This mask highlights regions that are statistically likely to be interacting with the juxta-membrane cortical actin as evidenced by the slower mobility of the membrane anchored binder (TractO) as compared to the non-binder (TractO*) in that pixel.

As the cortical actin is known to be dynamic and often it is built of fenestrae below the resolution limit (12, 21), the diffraction limited intensity values averaged over the whole duration might not have all the information about the actin dynamics. However, the average diffusion coefficient of the actin binders in a given pixel will be sensitive to the spatial heterogeneity of the cortical actin even if it cannot be resolved optically in the time averaged intensity maps (**Supplementary Figure S3**).

## Conclusion

The framework developed here establishes a practical route to functional mapping of the juxta-membrane actin cortex based on transport rather than intensity. In this context the weak binding regime is a good choice for two reasons. First, it reduces the likelihood that the probe will stabilize actin structures through prolonged binding and bundling as observed for experiments with Lifeact and F-Tractin (35–38). Second, it allows the probe to dynamically exchange between actin influenced environments during the acquisition. In this regime, the reporter’s mobility becomes a sensitive indicator of local actin influence, because even transient binding or repeated encounters with actin associated constraints slow down lateral mobility.

The two-channel design also generalizes naturally to other questions where a spatially heterogeneous regulator is suspected to shape membrane dynamics. For example, one can substitute the non-binder channel with a receptor, lipid probe, or signaling component and ask how its diffusion landscape covaries with the actin readout under perturbations that remodel the cortex. More broadly, the ability to compute transport-defined occupancy maps provides a quantitative bridge between qualitative actin imaging and models in which the cortex regulates reaction kinetics by shaping encounter rates, confinement, and effective search processes on the membrane.

## SUPPORTING MATERIAL

Supplementary Figures S1-3

Supplementary Movie 1

## ACKNOWLEDGMENTS

We thank all the members of the Mayor laboratory, in particular Thomas van Zanten and Bhagyashri Mahajan for discussions, suggestions and feedback on this project. We thank Amit Cherian and Greeshma Pradeep S for help with optical setups at NCBS. ST acknowledges doctoral fellowship support from National Centre for Biological Sciences, Tata Institute for fundamental Research (NCBS-TIFR).

## AUTHOR CONTRIBUTIONS

Conceptualization: ST, SM; Data Curation: ST; Formal Analysis: ST; Funding acquisition: SM; Methodology: ST, PK, TW, SM; Investigation: ST, PK; Visualization: ST; Project administration: SM; Software: ST, TW; Supervision: SM; Writing – original draft preparation: ST, SM; Writing – review & editing: ST, SM

## DECLARATION OF INTERESTS

Competing interests: Authors declare that they have no competing interests.

## FUNDING

Funding: SM acknowledges support from Department of Biotechnology – Wellcome Trust India Alliance Margadarshi Fellowship (IA/M/15/1/502018) and Leverhulme Trust, UK (LIP-2021-017). SM acknowledges the Department of Atomic Energy, India (under Project No. RTI 4006) and JC Bose National Fellowship (JBR/2021/000014). TW gratefully acknowledges funding by the Ministry of Education of Singapore (MOE-T2EP30223-0042)

---

**Figure S1: Optimized Fluorophore Configuration configuration for 2 channel ImFcs a)** Graph shows excitation and emission spectra for the two fluorophores where mStayGold is excited with the 488nm and the JF-657 dye is excited with the 640nm laser. Note minimal excitation and minimal emission cross talk. The emission filters used for collection after the beam splitter (gray line) are also highlighted. The spectra were generated using the spectra-viewer tool from https://fpbase.org. **b)** Normalized intensity traces of the C1 –mStayGold (green) and C2 – JF657 (blue) C2-Binder (blue points) during the acquisition process. Intensity trace of an eGFP labelled tracer from a different experiment with half the laser illumination plotted for comparison for photostability analysis (grey).

**Figure S2: mStayGold-TractO dynamics** This montage contains snapshots from a 4-minute long ImFCS acquisition, with frames acquired continuously every 2ms. The ∼120,000 frame movie was binned every ∼200 frames to generate intensity images that capture the dynamic nature of the binder intensity, Here snapshots from the binned movie are shown at 20 second intervals. The full movie is in the supplementary materials. The highlighted squares are a guide to the eye, to see regions with clear remodelling of actin intensity.

**Figure S3: Time averaged intensity v/s diffusion difference map a)** Representative diffusion difference map of the mStayGold-TractO in channel 1 and the HaloTag-JF657-TractO* in channel 2. **b)** Intensity ratio map, of the Channel 1 mStayGold-TractO intensity map divided by the Channel 2 HaloTag-JF657-TractO* intensity map. c) Scatter plot depicts the difference in diffusion coefficients from corresponding pixels from panel a plotted against the intensity ratio of the pixels from panel b.

**Supplementary Movie 1: mStayGold-TractO dynamics** Movie of the mStayGold-TractO construct described in the main text, imaged every ∼2ms in TIRF for 4minutes. The ∼120,000 frame movie was binned every ∼200 frames to generate intensity images that capture the dynamic nature of the binder intensity.

## Notes

### Competing Interest Statement

The authors have declared no competing interest.

