## Supplementary Material for "Two-colour Imaging Fluorescence Correlation Spectroscopy (ImFCS) probes the influence of the juxta-membrane actin-cortex"

FIGURE S1

(a)

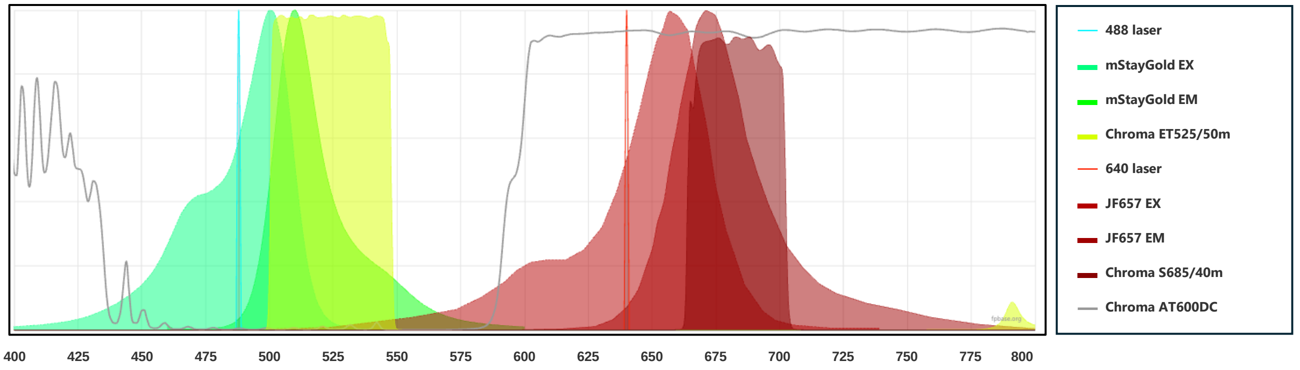


(b)

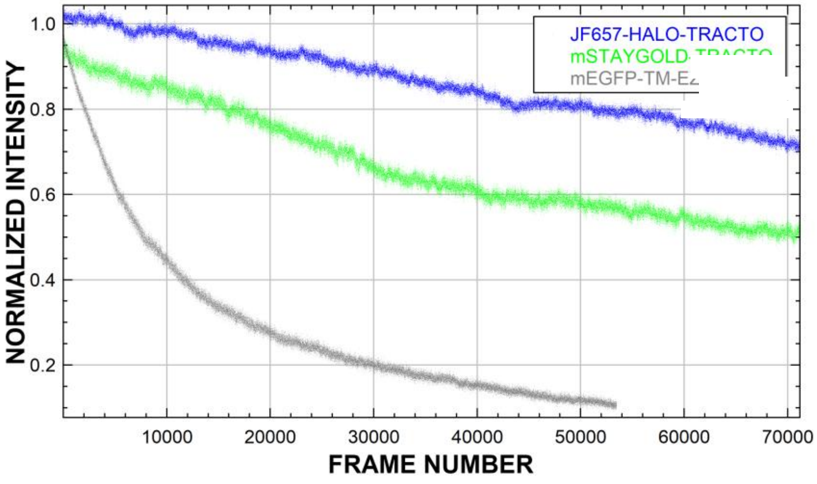


**Figure S1: Optimized Fluorophore Configuration configuration for 2 channel ImFcs**

**a)** Graph shows excitation and emission spectra for the two fluorophores where mStayGold is excited with the 488nm and the JF-657 dye is excited with the 640nm laser. Note minimal excitation and minimal emission cross talk. The emission filters used for collection after the beam splitter (gray line) are also highlighted. The spectra were generated using the spectra-viewer tool from fpbase.org. **b)** Normalized intensity traces of the C1 –mStayGold (green) and C2 – JF657 (blue) C2- Binder (blue points) during the acquisition process. Intensity trace of an eGFP labelled tracer from a different experiment with half the laser illumination plotted for comparison for photostability analysis (grey)

FIGURE S2

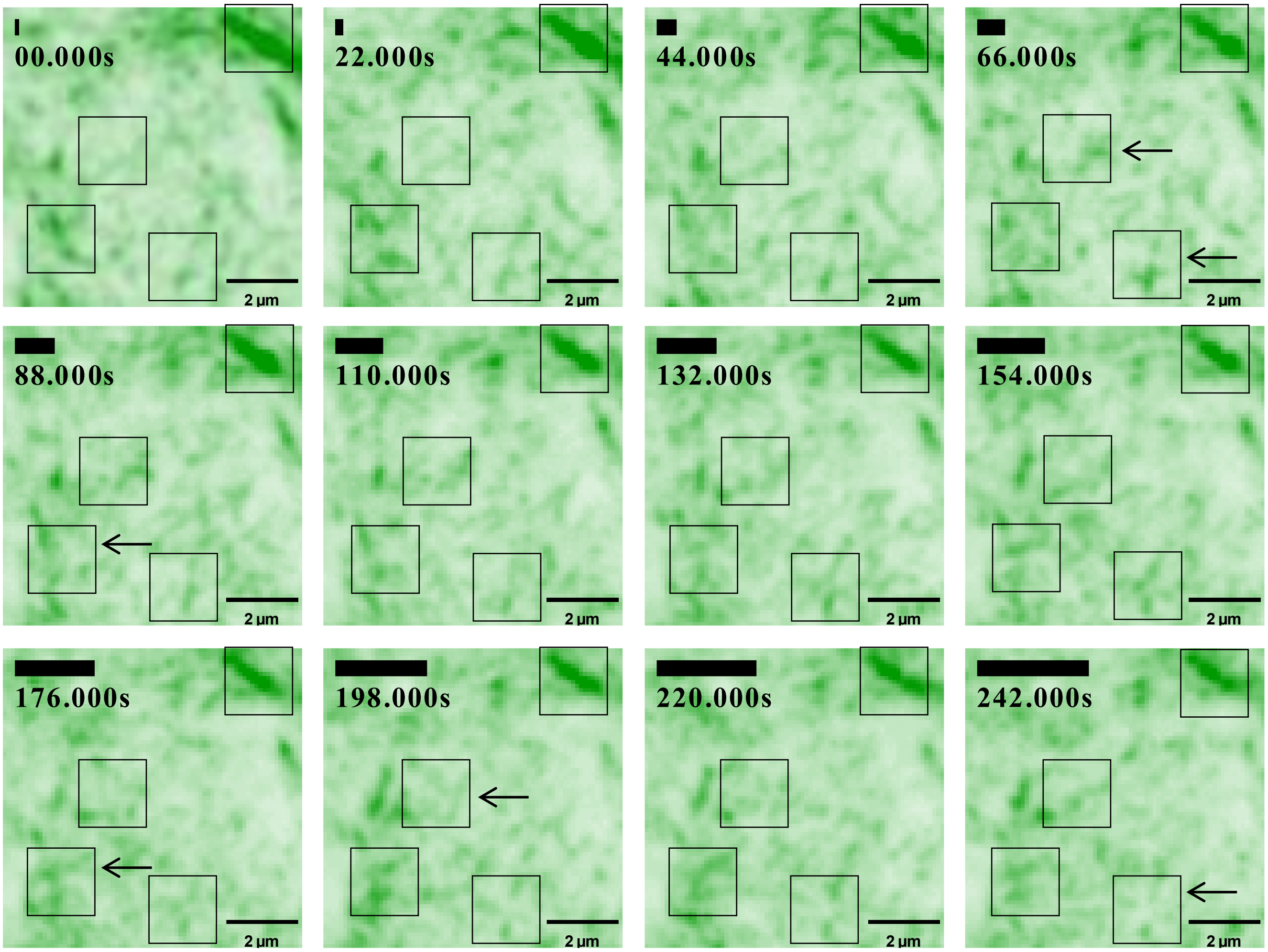


**Figure S2: mStayGold-TractO dynamics**

This montage contains snapshots from a 4-minute long ImFCS acquisition, with frames acquired continuously every 2ms. The ~120,000 frame movie was binned every ~200 frames to generate intensity images that capture the dynamic nature of the binder intensity, Here snapshots from the binned movie are shown at 20 second intervals. The full movie is in the supplementary materials. The highlighted squares are a guide to the eye, to see regions with clear remodelling of actin intensity

FIGURE S3

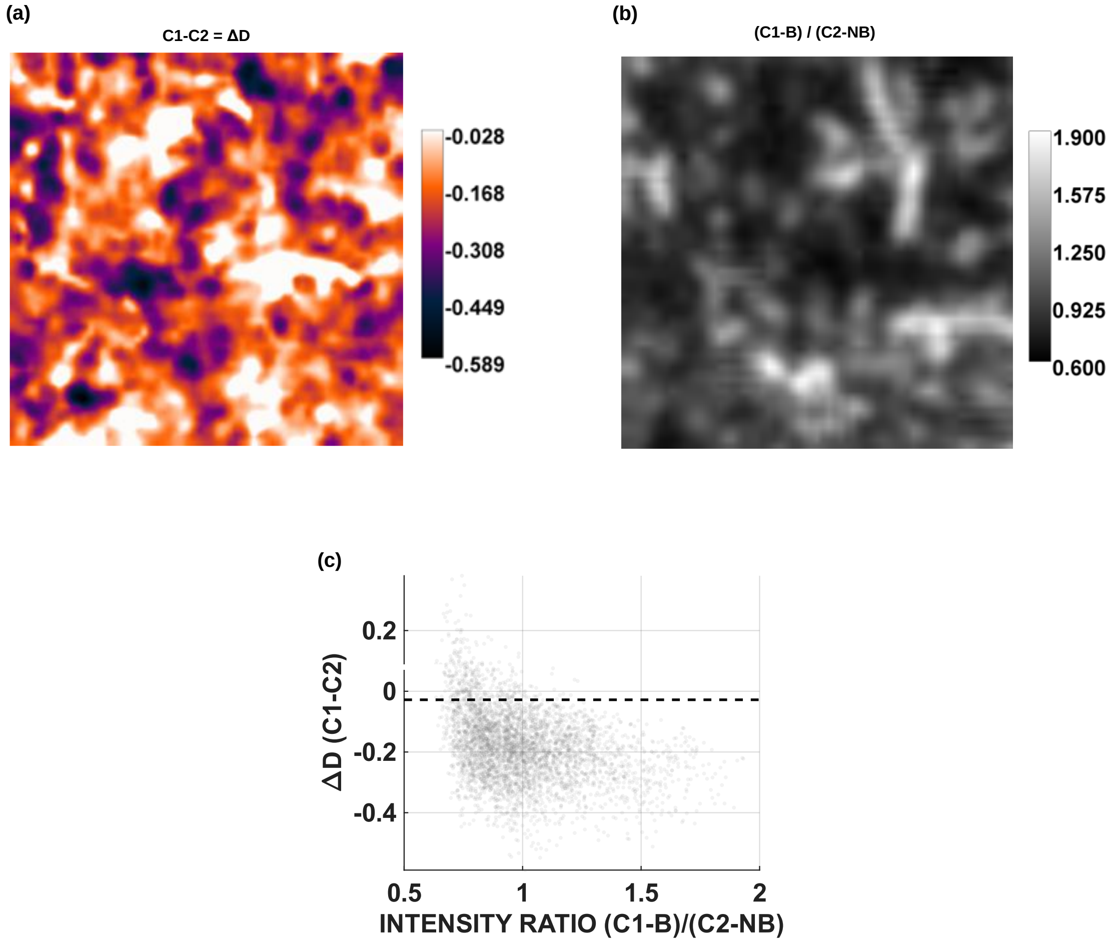


**Figure S3: Time averaged intensity v/s diffusion difference map**

**a)** Representative diffusion difference map of the mStayGold-TractO in channel 1 and the HaloTag-JF657-TractO* in channel 2. **b)** Intensity ratio map, of the Channel 1 mStayGold-TractO intensity map divided by the Channel 2 HaloTag-JF657-TractO* intensity map. c) Scatter plot depicts the difference in diffusion coefficients from corresponding pixels from panel a plotted against the intensity ratio of the pixels from panel b
